# Factors Affecting Germination of a Dominant Salt Marsh Species are Context-Dependent: Implications for Coastal Seed-Based Restoration

**DOI:** 10.64898/2026.04.23.720394

**Authors:** Bethany J. Lee, Kerstin Wasson, Monique C. Fountain, Rikke Jeppesen, Zeanna Graves, Wesley P. Moore, Margaret Zimmer, Anna E. Braswell

## Abstract

Restoration of coastal salt marshes is often limited by their capability to revegetate, either through natural recruitment or active planting methods. Despite the critical need for efficient revegetation methods, direct seeding remains an underrepresented approach in coastal wetland restoration. Additionally, tidal inundation poses special challenges for coastal Seed-Based Restoration (SBR) relative to terrestrial habitats, with tides displacing seeds from the marsh platform. To determine factors potentially influencing successful establishment in coastal wetlands, we conducted a series of greenhouse and lab experiments with *Salicornia pacifica* (pickleweed), the dominant plant in California marshes. We determined pickleweed seed viability using standardized germination tests. Additionally, we tested factors that influence pickleweed seed viability and germination rates, such as soil moisture, soil type, and sowing depth. We found that pickleweed seeds had an average viability of 22.5%, which increased with larger-sized seeds. We also determined that the most effective dormancy-breaking pretreatments varied by soil type: a one-day cold stratification in freshwater maximized germination in benign soils, whereas a seven-day cold stratification in saltwater maximized germination in stressful soils. Finally, we determined that the optimal conditions for sowing seeds are surface sowing under moderate moisture. Creating conditions to maximize viability and germination is crucial to ensure the greatest chance of successful revegetation post restoration. Our study, which sequentially tested factors affecting different phases of the early life history of a dominant foundation species, can inform SBR for other coastal plants. This approach can help coastal land managers successfully implement SBR for habitat restoration.

## Introduction

Coastal salt marshes provide critical ecosystem services such as carbon sequestration, storage of toxic metals, and sources of organic matter and nutrients (Zedler and Kercher, 2005), and are increasingly threatened by sea level rise and coastal development, with extensive losses already documented (Valiela *et al*., 2018). Land managers and municipalities use coastal wetland restoration to combat marsh loss and preserve wetlands (Donatelli *et al*., 2018; Billah *et al*., 2022). Coastal restoration projects employ different methodologies for revegetation, including planting nursery-grown plugs, transplanting vegetation from adjacent marshes, or relying on natural recruitment (Billah *et al*., 2022). While natural recruitment can be a slow process, active planting methods have demonstrated rapid revegetation success but are often resource-intensive and costly to implement.

Seed-Based Restoration (SBR) is a widely used tool for restoring terrestrial systems (Martin and Wilsey, 2006; Vitt *et al*., 2022). In this approach, seeds from a single species or a blend of multiple species are sown across an area to enhance biodiversity, add cover to bare areas, or facilitate future vegetation establishment (Kardol *et al*., 2008; Kettenring and Tarsa, 2020). Seed additions have been successful in grasslands (Martin and Wilsey, 2006), woodland forests (Kaul *et al*., 2023), and prairies (Kardol *et al*., 2008). In coastal systems, most documented uses of seed additions are for seagrasses. For example, the Chesapeake Bay has implemented large-scale seed-based restoration of *Zostera marina*, which has successfully revegetated over 3,500 ha of seagrass meadows (Orth *et al*., 2020). Coastal wetlands could also benefit from seed addition, but there are few documented applications. Limitations are largely due to wave action displacing seeds placed on lower marsh platforms and a lack of knowledge of coastal seed biology. As sea levels rise, marshes are expected to migrate to higher elevations, with a gradual transition of ecotones from upland terrestrial systems to high marsh (Wasson, Woolfolk and Fresquez, 2013). These high marsh areas could greatly benefit from the application of SBR to acclimate and prepare the ecosystem for the effects of sea level rise. Other areas that would be enhanced from SBR efforts include large construction restoration projects, such as large-scale sediment additions, where the resulting landscape begins as bare sediment that needs to be revegetated.

To maximize the success of SBR for coastal marshes, each step of the seed addition process must be carefully evaluated—from seed sourcing and storage to pre-treatment and sowing conditions (Table 1). Seed sourcing should consider localized adaptations to salinity gradients, tidal inundation, and sediment conditions, as seed provenance can strongly influence viability and physiological thresholds (Vitt *et al*., 2022). Once collected, storage practices, such as maintaining adequate humidity or temperature, are essential for preserving seed vigor, especially for species with short-lived seeds (De Vitis *et al*., 2020). Appropriate pretreatments, such as stratification, scarification, or light exposure, may be required to overcome dormancy and initiate germination during favorable tidal or moisture windows (Koornneef, Bentsink and Hilhorst, 2002). The sowing environment determines the success of the germinant, as factors such as microtopography, moisture availability, and light availability influence both germination and early seedling establishment (Lamichhane *et al*., 2018). Finally, the transition from germinant to established seedling represents a critical bottleneck in restoration, where coastal stressors such as desiccation, inundation, wave action, or competition can determine long-term persistence (Zhao *et al*., 2023). Because of the stressful nature of certain restoration methods (e.g. bare sediment, lack of present organics) it is crucial to understand the stages of sowing and seedling establishment under standard, non-stressed conditions, as well as under environments of high stress. Site-specific stressors may lead to different establishment outcomes and restoration trajectories (Valliere *et al*., 2022).

**Table 1.** Stages of seed additions that are addressed in our study and that are critical to restoration success.

| **Critical Stages of Seed Additions** | **A.**<br>**<u>Sourcing of Seeds</u>** | **B.**<br>**<u>Storage</u>** | **C.**<br>**<u>Pretreatment of Seeds</u>** | **D.**<br>**<u>Sowing Seeds</u>** | **E.**<br>**<u>Seedling Establishment</u>** |
| --- | --- | --- | --- | --- | --- |
| **Environmental Influences on Establishment Success** | Maternal environmental or genetic conditions affecting seed production, viability, and embryonic fill | Temperature and humidity<br>Duration of storage | Scarification<br>Stratification<br>Photoperiod Exposure | Soil compaction<br>Soil salinity<br>Soil moisture<br>Nutrient availability | Precipitation<br>Granivory<br>Competition (Inter- and Intraspecific) |
| **Manager Intervention to Enhance Success** | Choose seeds from high viability sources | Store seeds in conditions conducive to maintaining viability | Treat seeds to break dormancy and maximize germination | Sow seeds into the soil in favorable environmental conditions | Maintain moisture to prevent desiccation and prevent early seedling death |

Along the western coast of North America, pickleweed (*Salicornia pacifica*) occurs in salt marshes, often as the dominant species in the Mediterranean climate at the mid- to southern end of its range (Kadereit *et al*., 2007). Pickleweed is a well-known, hardy species, and several studies have investigated factors shaping its distribution and natural recruitment (Skougard and Brotherson, 1979; Zedler, 1983; El-Nwehy *et al*., 2020). One coastal study that examined establishment outside of natural recruitment focused on spreading broken stems as vegetative material, but few have reported using direct seeding methodologies (Miles *et al*., 2015). Average viability and germination rates of the Salicornia genus significantly vary among their species and on environmental factors like salinity or light exposure *(cite calone 2020 and Khan 2000 and Gunaskara 2025)*. For example, *Salicornia europea* reached a peak germination of 43% only when exposed to 860mM salt concentrations in water at 5-15°C (Ungar, 1977). Pickleweed establishes passively from tide-dispersed seeds at California restoration sites at frequently inundated, low elevations (Zedler, Morzaria-Luna and Ward, 2003; Williams and Faber, 2004). This passive approach, relying on tidal dispersal, may not result in abundant pickleweed cover or rapid colonization on higher marsh platforms that are less frequently inundated, or on platforms under increasing drought frequency and intensity associated with climate change. SBR could be used to increase pickleweed cover under such conditions; however, more research is required to maximize success.

This study focused on several critical stages of SBR for pickleweed and was specifically motivated by the potential to implement SBR at a high-elevation marsh restoration site in the Elkhorn Slough estuary in central California, USA. First, we examined seed viability and its potential correlation with seed size through standardized germination trials in the lab. To determine whether some source populations provide a higher percentage of viable seeds than others—and thus would be most effective for restoration—we tested which sources within Elkhorn Slough produced higher-quality seeds. We also examined whether larger seeds, potentially indicative of more developed embryos (Baskin & Baskin 2022), had higher germination rates. Next, we evaluated various pre-treatment methods to enhance germination and break dormancy. This included a full-factorial stratification experiment, along with cold saltwater soaking to simulate overwintering and tidal exposure. We also investigated the effects of salinity and moisture on germination success, as desiccation is a common barrier to seedling establishment. Since pickleweed is a halophyte, we tested whether a minimum salinity threshold is required for germination, and conducted a submersion experiment to determine optimal moisture levels. Finally, we assessed how sowing depth influences germination outcomes. The approach we implemented, sequentially testing critical factors involved in different stages of SBR (Table 1), can serve as a model for SBR in other systems.

## Methods

### Focal System - Hester Marsh

Our greenhouse experiments were motivated by the potential for implementing SBR at a restoration site, Hester Marsh in Elkhorn Slough. This estuary is located about 50 miles south of San Francisco on the Central Coast of California, USA, in a region dominated by increasing climate fluctuations and drought stress (Deitch, Sapundjieff and Feirer, 2017). Previously, anthropogenic diking led to ground subsidence that converted the marsh into a mudflat. Soil added to Hester Marsh restored elevation levels to approximately half a meter higher than the marsh originally was. The soil added to the marsh plain came from a nearby flood reduction project and the lower soil layers of an adjacent fallow farm. Both soil sources had a low organic matter content of around 2-4%. The soil was primarily a sand/silt mix that was heavily compacted on the restoration site as a result of the construction equipment used to move the soil (Wasson et al. 2025). The goal of this project was not only to restore the marsh but also to build it high enough to be climate-ready, able to withstand 50 cm of sea-level rise. Hester’s first phase of construction finished in 2018, when it was opened to the tides. Natural recruitment by seeds arriving on tides resulted in only 30% cover (primarily by pickleweed) seven years after construction of this area (Wasson et al. 2025), pointing to a potential benefit to active restoration with SBR. We initially tested various SBR methods in small-scale field experiments at Hester Marsh, but there was no evidence of seed germination or establishment. Therefore, the goal of the current greenhouse experiments was to inform SBR efforts at Hester.

### Baseline Viability

To determine the baseline viability of the pickleweed seeds (Table 1A), we conducted germination tests using the standardized petri plate method (Baskin and Baskin, 2014). The seeds used were sourced from 9 different harvests across Elkhorn Slough, collected in October 2023. The 9 sources varied across different elevations, different inundation frequencies, distances from the shoreline, and other environmental stressors (Figure S1). Within each source, the whole site was not evenly represented; only multiple high-elevation plants within each site were collected to compare high marsh sites across the Slough. High-elevation for our model site was approximately 2 meters above sea-level. After seed collection, seeds were dried out for one week, then filtered for chaff, and stored in the fridge. Seeds were soaked for one week in freshwater in a 12°C refrigerator. Once stratification was completed, seeds were placed on moist germination paper, in petri plates under a grow light for 8 hours dark - 16 hours light for 21 days. There were 50 seeds per petri plate with 3 replicates of each plate (Baskin and Baskin, 2014). With 10 sources, that led to 30 total plates. Seeds were monitored every other day, with additional moisture added around the germination paper to prevent the seeds from drying out. Seeds were scored as “germinated” when the radicle emerged from the seed (Baskin and Baskin, 2014).

We conducted all statistical analyses for this section and the following in R version 4.2.3 (R Core Team 2023). We used a generalized linear model (GLM) with a binomial distribution to test for differences in germination proportions among seed sources. We then performed Tukey-adjusted post-hoc pairwise comparisons to identify which sources differed significantly in germination success.

### Seed Sizing

Using the same replicates from the viability experiment, which included ten seed sources, each seed was measured under a microscope to calculate an average seed size for each petri dish (Baskin and Baskin, 2014). This was repeated across all 30 treatments listed above, yielding measurements for 1,500 seeds. The average seed size per replicate was then paired with the corresponding viability test results for that petri dish.

Three generalized linear models (GLMs) were run to assess sizing and viability: (1) A model of germination percentage as a function of average seed size while including site as a factor, (2) a model of germination percentage as a function of average seed size with site removed as a factor, and (3) a model of germination percentage as a function of seed size with a categorical inundation frequency (frequent or infrequent inundation) of the source marsh as an alternative variable for site. Frequent inundation source plants experienced tidal inundation action once per day, while infrequently inundated sources were exposed to less than daily tidal inundation (typically only a few times a year). Germination models were fit with a quasibinomial distribution, as model diagnostics using the DHARMa package found mild overdispersion (dispersion = 1.68).

### Stratification

To test the efficacy of different stratification methods and the variation in germination across soil types (Table 1C), we conducted a greenhouse experiment at the University of Florida in Gainesville, Florida. Seeds were sown in two soil types: 1) Hester mimic soil, with high clay content and very low organic content, and 2) potting soil, providing a contrast with higher organic content than the restoration site. We conducted a full factorial experiment: pickleweed seeds were either soaked in freshwater (0 ppt) or saltwater (30 ppt), at room temperature (20°C) or cold water (12°C), for either 3 hours, 1 day, or 7 days. Soaking times were selected to mimic potential time frames the seeds remained in the tides before returning to the marsh platform. The seeds were then sown into either Hester mimic soil or into potting soil immediately after the soaking ended. Each soil type included a control group of seeds that were sown directly without any soaking. There were six replications for each combination of the 24 different possible treatments (144 total planted pots; Figure S2). Each of the 144 planting pots (∼3.2 liters) contained 50 seeds. After sowing, seeds were monitored daily for one month; the first germination occurred on day 3, and the last on day 17 in March 2023.

We conducted additional soil tests were conducted on six samples per soil treatment, collected from the center of randomly selected pots, to compare salinity, organic matter, and particle size between the potting soil and the Hester mimic soil (Figure S3). Soil cores were taken using a push corer and stored in sealed bags until processing later that day. Each sample was weighed, dried in a drying oven, and re-weighed to determine gravimetric water content. Dried samples were then ground and homogenized before subsampling for organic matter and salinity analyses. Organic matter was quantified using a loss-on-ignition procedure, in which 10 g subsamples were combusted in a muffle furnace at 500°C for four hours and then re-weighed to determine percent organic matter. For salinity, a 2.5 g subsample was diluted with deionized water at a 1:10 ratio, and specific conductivity and salinity (ppt) were measured using a YSI probe (Xylem XA00088-02). Particle size distribution and corresponding soil textural class were assessed using a particle size analyzer at the University of California, Santa Cruz.

Germination responses were analyzed using GLMs. Germination percentages were modeled with a binomial error distribution using the number of germinated seeds (grs) and the approximate number of ungerminated seeds (50 - grs) from the GerminaR output (Lozano□Isla, Benites□Alfaro and Pompelli, 2019). Mean germination time was modeled with a gamma distribution to account for its positive, right-skewed distribution. Both models included soil type, soak temperature, soak time, and salinity soak as fixed factors, with an interaction between soak time and salinity soak. Post-hoc comparisons of soaking time and salinity were conducted using Tukey-adjusted pairwise tests on estimated marginal means (*emmeans*) to identify specific treatment differences.

### Effect of Soil Moisture and Water Salinity

To test the effect of soil moisture and salinity on germination percentage (Table 1D), and whether this differs by soil type, greenhouse experiments took place at a University of Florida greenhouse in Gainesville, Florida. Prior to seed addition, soils in the pots were watered either with fresh or saline water (30 ppt generated with InstantOcean), approximately 1 liter total to each treatment tray. After the initial addition of freshwater or saltwater, seeds were then sown after being soaked for 7 days in freshwater. Moisture levels were maintained at either high, medium, or low levels with daily freshwater additions and were monitored with moisture probes (ZENTRA Teros 12). High moisture levels were around 96% Volumetric Water Content (VWC), medium moisture levels were around 63% VWC, and low moisture levels were around 35% VWC. These reported moisture levels represent the average VWC immediately following daily watering in the greenhouse. VWC is the ratio of water volume to the total soil volume, expressed as a percentage. In addition to variations in moisture and salinity levels, the seeds were also sown in either hester mimic or potting soil. Each combination of treatments held approximately 0.1g of seed material, or around 146 ± 9 seeds per treatment. After seeds were sown, they were monitored, and germination was assessed daily for one month, with the first germination occurring on day 3, and the last occurring on day 23 in April 2023.

Germination percentage (number germinated out of the 146 total) was modeled using a beta-binomial GLM with a logit link (glmmTMB) to account for overdispersion. Soil type, salinity, and moisture were included, as well as all two-way and three-way interactions. Type II chi-square tests were used to assess the significance of main effects and interactions, residuals were evaluated with the DHARMa R package (Hartig, 2022), and *emmeans* were evaluated with Tukey-adjusted pairwise contrasts to compare treatment combinations. An additional model evaluated germination time using a Gamma GLM (log link) to account for the positive, right-skewed time responses. Similar Type II chi-square tests were done with DHARMa diagnostics, and *emmeans* were evaluated; all done on a log-Gaussian model.

### Submersion and Moisture

To determine the effect of soil moisture and seed submersion on germination (Table 1D), we conducted an experiment at the Elkhorn Slough National Estuarine Research Reserve (ESNERR). Twenty plastic cups (∼60 mL) were filled with potting soil to 1 cm below the rim and assigned to one of four soil moisture treatments: high (175 mL H□O), medium-high (125 mL), medium-low (100 mL), and low (60 mL). A fifth treatment consisted of seeds kept submerged in freshwater with no soil (“water-only”). Volumetric water content (VWC) was measured using a ProCheck Soil Moisture Meter and maintained as consistently as possible throughout the experiment. High-moisture treatments were maintained at 100% VWC, Medium-high at 50% VWC, medium-low at 25% VWC, and low at 10% VWC. Watering to maintain these VWC levels occurred during the monitoring days, which were approximately every three days. All seeds were soaked in freshwater for 1 week at 12°C prior to sowing. The soaked seeds were then lightly buried approximately 3mm deep in the soil. The cups were placed outdoors in a clear-lidded container under ambient conditions and monitored for 21 days, resulting in 7 monitoring dates occurring in July 2023. Germination percentage was analyzed using a quasibinomial generalized linear model (logit link) to account for significant overdispersion. Moisture treatment was included as a fixed factor, and model significance was evaluated using Type II Wald χ^2^ tests. Post-hoc pairwise contrasts were assessed with Tukey adjustment using the *emmeans* R package.

### Sowing Depth and Soil Type

To determine the effect of varying planting depth on germination (Table 1D), and how this varies with soil type, an ESNERR-based outdoor greenhouse experiment was conducted that compared Hester soil and the control potting soil at the depth at which seeds were sown. Hester soil was harvested from the top 12 cm of the site, sieved for seeds and large matter, and added to 266 mL plastic cups, with each soil type represented in 10 cups. Each category was evenly split between seed treatments - one representing surface seed dispersal and the other at 0.5 cm burial depth. Soil was filled and slightly compacted, 1 cm from the lip of each surface seed cup and 1.5 cm for the buried seed cups. Once seeds (n=50 per cup) were placed in the cups, 0.5 cm of soil was added over the seeds and lightly compacted down.All seeds were soaked in freshwater for 1 week at 12°C prior to sowing. Seeds were then sown into the soil with 100 mL of filtered fresh water. They were then kept saturated by adding fresh water as needed. The cups were kept outside, uncovered, during July 2023. The experiment was monitored for germination every few days for three weeks.

We used a GLM with a binomial error distribution to evaluate the effects of sowing depth (surface vs buried), soil treatment (Hester soil vs potting soil), and their interaction on germination outcomes. The response variable was the number of seeds that germinated out of 50, to model the probability of germination per seed, with 50 trials per cup.

## Results

### Baseline Viability

Seed viability was higher from sources that were infrequently inundated (p *=* 0.013) (Figure 1). Infrequently inundated sources had an average viability of 28.6% ± 2.27%, while frequently inundated sources averaged 7.11% ± 2.24%. Across all sources of pickleweed that were harvested, the average germination was around 22.5%. The seeds sourced from within the restoration site had the highest viability across all 10 sources, with individual plate percentages reaching as high as 60%, but averaging around 40% for Hester-sourced seeds. The adjacent, lower elevation site (source 8) had an average viability of around 2%, with only 4 seeds germinating on average across all test dates. The sources that were most frequently inundated had the lowest germination rates (Sources 4, 7, and 8).

**Figure 1.**
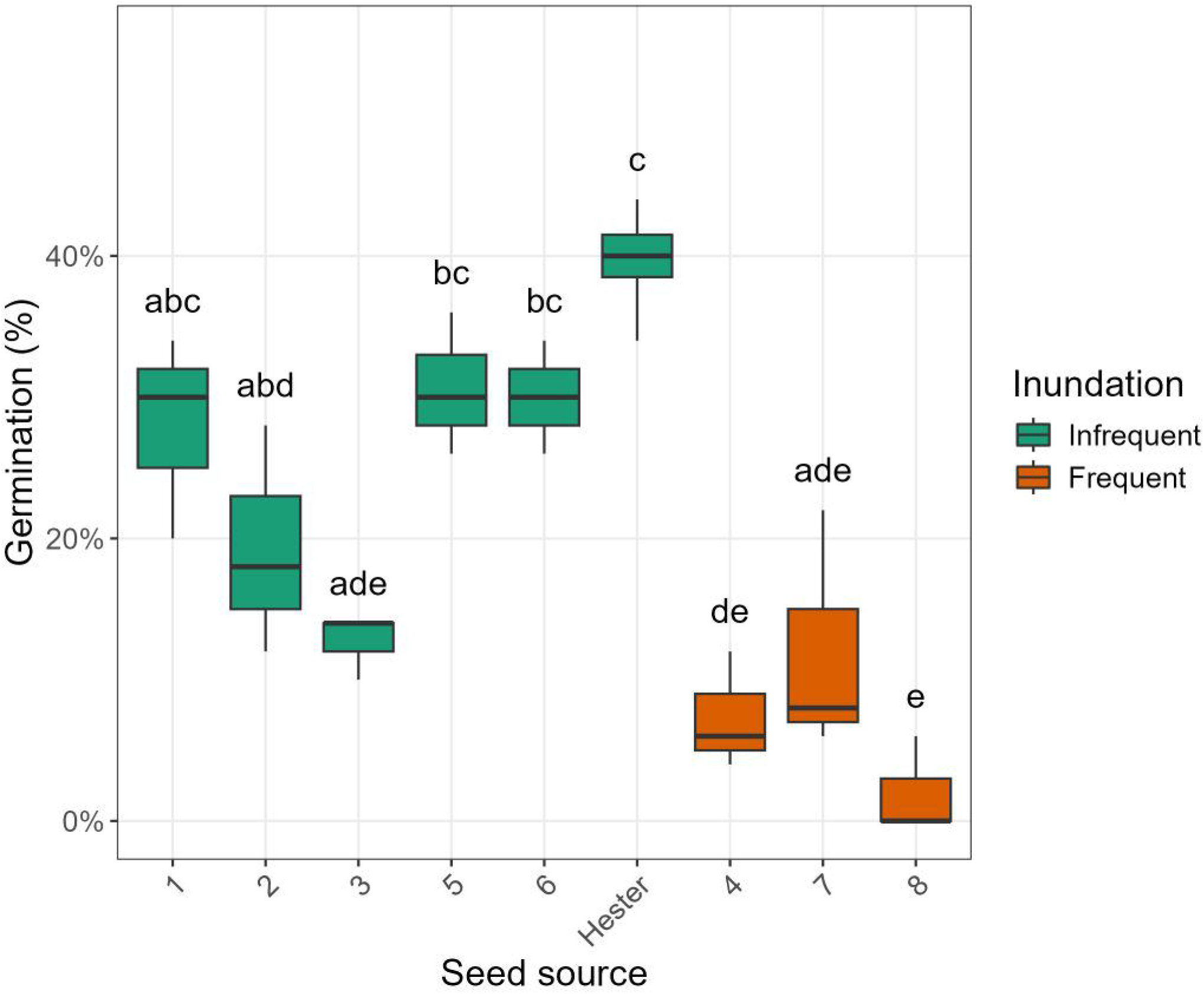
Germination percentage of pickleweed seed viability across the 9 sampled sources (numbers on x-axis represent site numbers). Each box represents the interquartile range with the median indicated as the middle line. The whiskers extend to 1.5 times the interquartile range, with no outliers present. To indicate the statistically supported differences among sources, Tukey-adjusted pairwise comparisons of the estimated marginal means were converted for a letter display. Groups that do not share a letter differ significantly at □ = 0.05, whereas sources sharing the same letter are not significantly different.

### Seed Sizing

Seed size varied significantly by source site, with the largest seeds originating from Hester (Figure S4). The GLM assessing germination as a function of average seed size included site as a factor and revealed a significant positive effect of seed size on germination probability (p = 0.013, z = 2.48, SE = 2.88), indicating that larger seeds were more likely to germinate (Figure S5). This relationship between seed size and germination probability does not hold for within-site analysis.

Using a quasi-binomial GLM to examine the effects of source inundation frequency and size on germination, we found that seeds from sites with frequent inundation had significantly lower germination than seeds from those with infrequent inundation. No significant relationship was found between seed source inundation frequency and seed size. An odds ratio of around 0.51 indicated a 49% lower odds of germination of seeds from frequent inundation sites.

### Stratification

Soak time had a significant effect on germination percent (Table 2). Across all soil and salinity treatments, germination increased with longer soak durations, with the 7-day soak resulting in the highest overall germination (27.4%). Soil type also significantly influenced germination (Table 2), with seeds in potting soil germinating 5 times more often than those in the Hester mimic soil (Figure 2). The control (unsoaked) treatment reduced germination, especially in the Hester Mimic soil, where no seeds germinated. The interaction between soaking temperature and salinity was significant for cold-water soaks (p < 2e-16), resulting in higher germination when compared to room-temperature soaks.

**Table 2.**
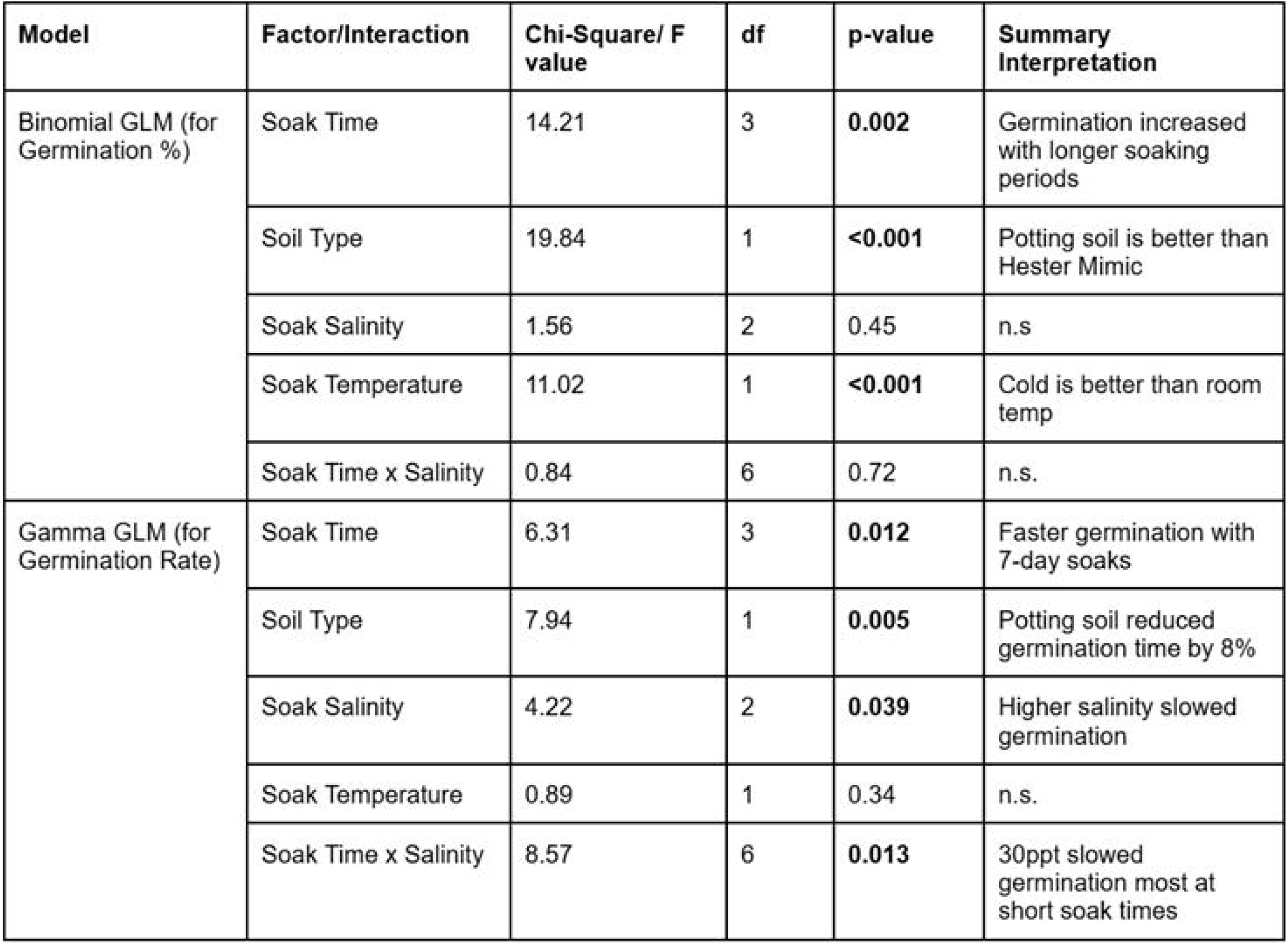
Results of generalized linear models testing the effects of seed soaking treatments and soil type on pickleweed seed germination. n.s. stands for non-significant.

| Model | Factor/Interaction | Chi-Square/ F value | df | p-value | Summary Interpretation |
| --- | --- | --- | --- | --- | --- |
| Binomial GLM (for Germination %) | Soak Time | 14.21 | 3 | **0.002** | Germination increased with longer soaking periods |
|  | Soil Type | 19.84 | 1 | **<0.001** | Potting soil is better than Hester Mimic |
|  | Soak Salinity | 1.56 | 2 | 0.45 | n.s |
|  | Soak Temperature | 11.02 | 1 | **<0.001** | Cold is better than room temp |
|  | Soak Time x Salinity | 0.84 | 6 | 0.72 | n.s. |
| Gamma GLM (for Germination Rate) | Soak Time | 6.31 | 3 | **0.012** | Faster germination with 7-day soaks |
|  | Soil Type | 7.94 | 1 | **0.005** | Potting soil reduced germination time by 8% |
|  | Soak Salinity | 4.22 | 2 | **0.039** | Higher salinity slowed germination |
|  | Soak Temperature | 0.89 | 1 | 0.34 | n.s. |
|  | Soak Time x Salinity | 8.57 | 6 | **0.013** | 30ppt slowed germination most at short soak times |

**Figure 2.**
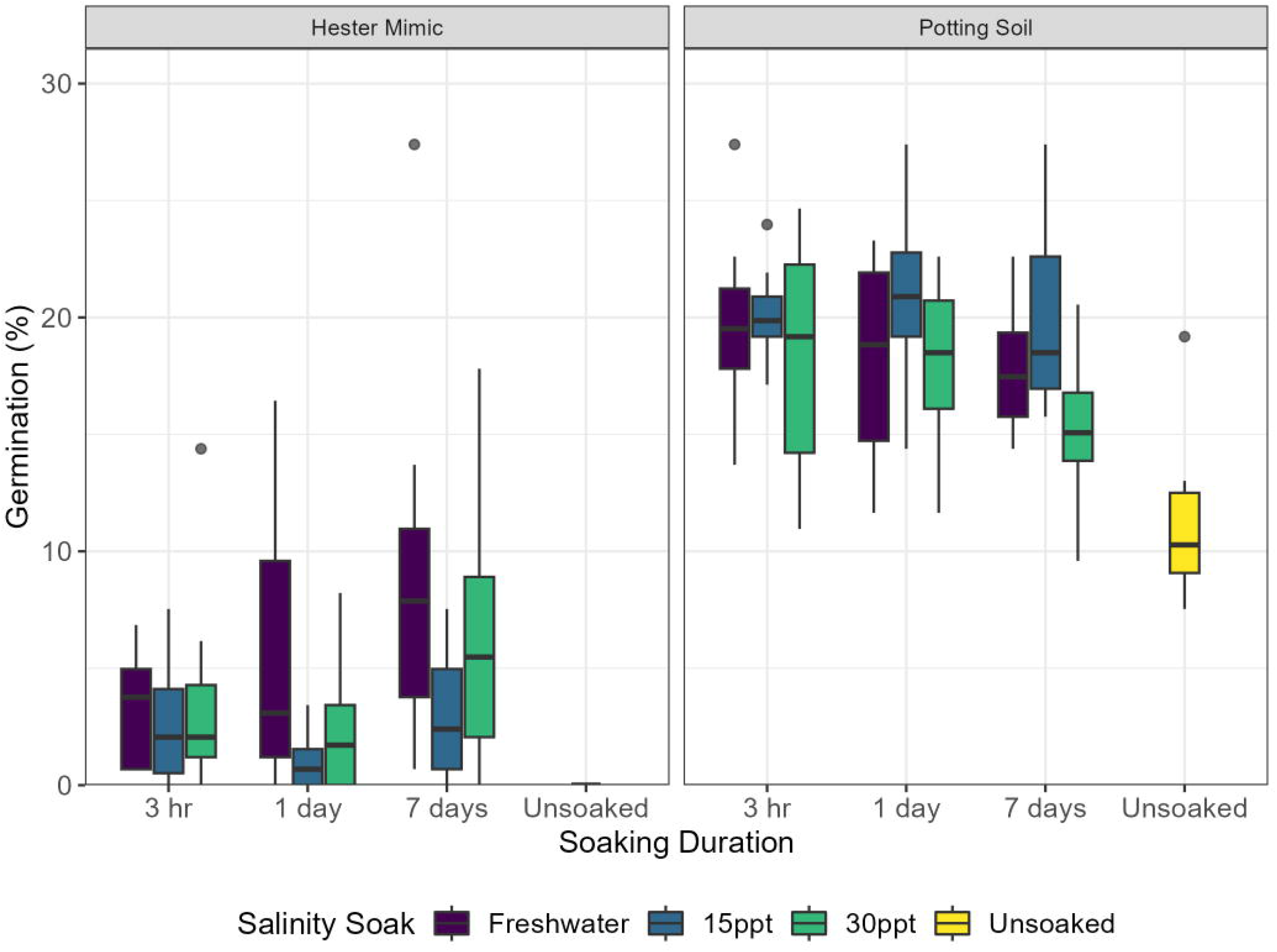
Differences in the number of germinated seeds depending on soil treatment after different stratification treatments of pickleweed seeds. Each box represents the interquartile range with the median indicated as the middle line. Whiskers extend to 1.5 times the interquartile range, with outliers presented. Hester mimic soil (left) resulted in overall lower germination than potting soil (right). Unsoaked seeds did not germinate in the hester mimic soil, and had the lowest overall germination of all the treatments for the potting soil.

Soak time, soil type, and salinity were all significant predictors of mean germination rate. The 7-day soak treatment led to faster germination, meaning around 3 days to germinate, whereas high salinity significantly delayed germination by roughly 21%, or around one day (Table 2). Potting soil reduced the mean germination time by about 8% compared to the Hester Mimic soil (Table 2). A significant interaction between soak time and salinity (Table 2) indicated that the slowing effect of high salinity was strongest at short soak durations. No significant effects of soaking temperature on germination timing were detected.

### Effect of Soil Moisture and Water Salinity

Results from the moisture and salinity experiment were highly dependent on the soil in which the seeds were sown (Figure 3). Likelihood-ratio comparisons confirmed a three-way interaction between soil type, salinity, and moisture (X^2^ = 8.86, p = 0.0119), and the sets of two-way interactions (X^2^ = 30.80, p < 0.005). The type II test in the full model showed significant effects of soil (p < 0.001), salinity (p = 0.006), and significant Soil X Salinity, Soil X Moisture, Salinity X Moisture, and Soil X Salinity X Moisture interactions (all p < 0.01). The post-hoc test showed the greatest amount of germination occurred in potting soil under saltwater treatments (0.25-0.28), and the lowest amount of germination in Hester mimic soil under the saltwater, low moisture treatments (0.07).

**Figure 3.**
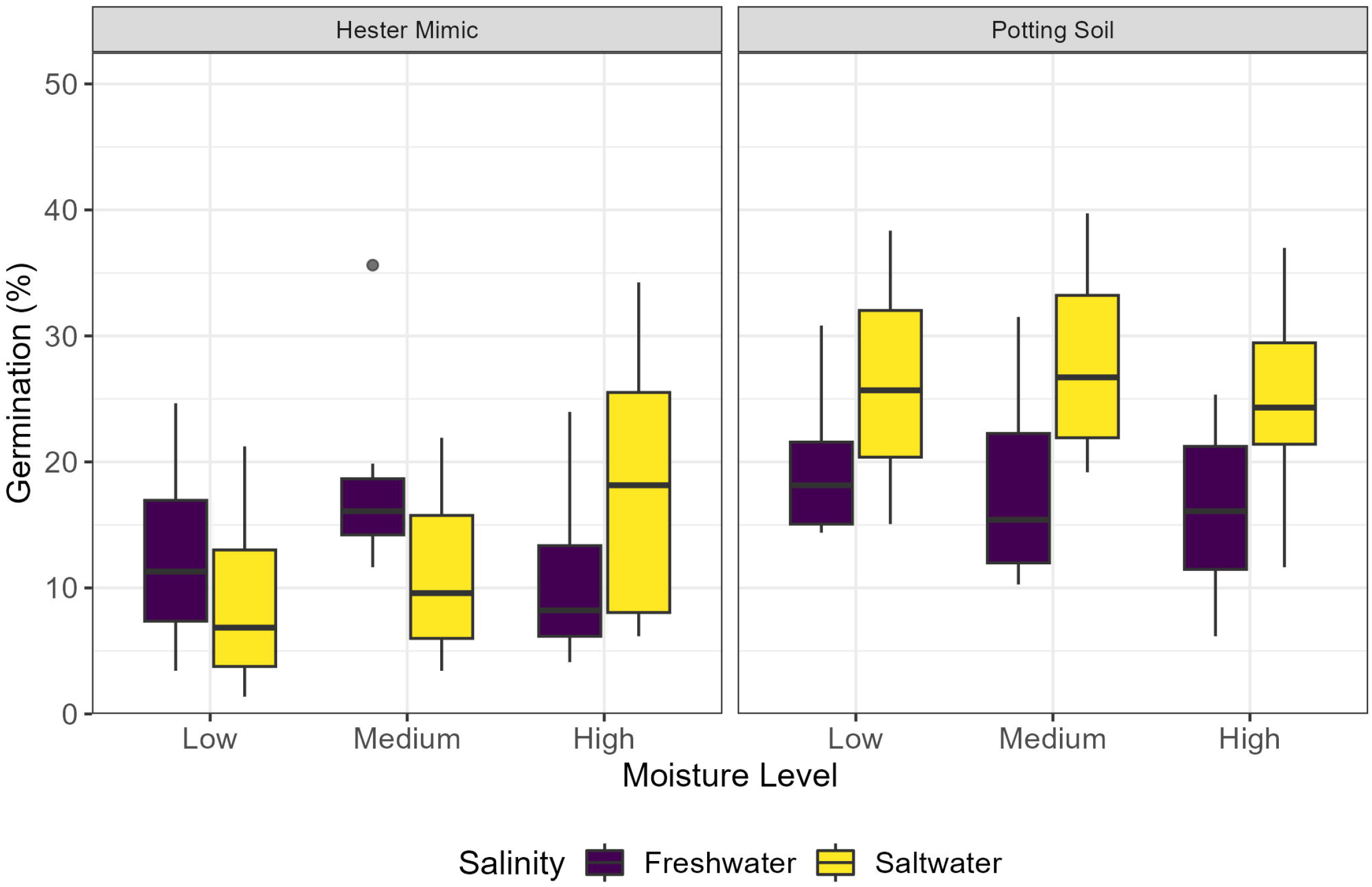
Differences in germination time and the number of germinated seeds by soil substrate and moisture treatments of pickleweed seeds. Each box represents the interquartile range with the median indicated as the middle line. Whiskers extend to 1.5 times the interquartile range, with a single outlier. High moisture levels were around 96% VWC, medium moisture levels were around 63% VWC, and low moisture levels were around 35% VWC.

The model for germination time, which used a Gamma (log) GLM, showed no three-way interaction among the variables. Our type II test showed significant effects of soil (X^2^ = 19.34, p < 0.01), salinity (X^2^ = 4.65, p = 0.031), and an interaction between salinity and moisture (X^2^ = 6.48, p = 0.039). The predicted means ranged from around 10 days (potting soil, saltwater, low moisture treatment) to around 14 days (Hester mimic, saltwater, high moisture, Figure S7).

### Submersion and Moisture Treatments

Germination differed significantly among the five moisture treatments (χ^2^ = 206.53, p < 0.001; Figure 4). Seeds that remained submerged in water exhibited the highest germination probability (estimated 0.57, 95% CI: 0.49–0.64, Figure S8). In contrast, all four soil-based moisture treatments produced similarly low germination probabilities (p < 0.10), and no significant differences were detected among them (all pairwise p > 0.05).

**Figure 4.**
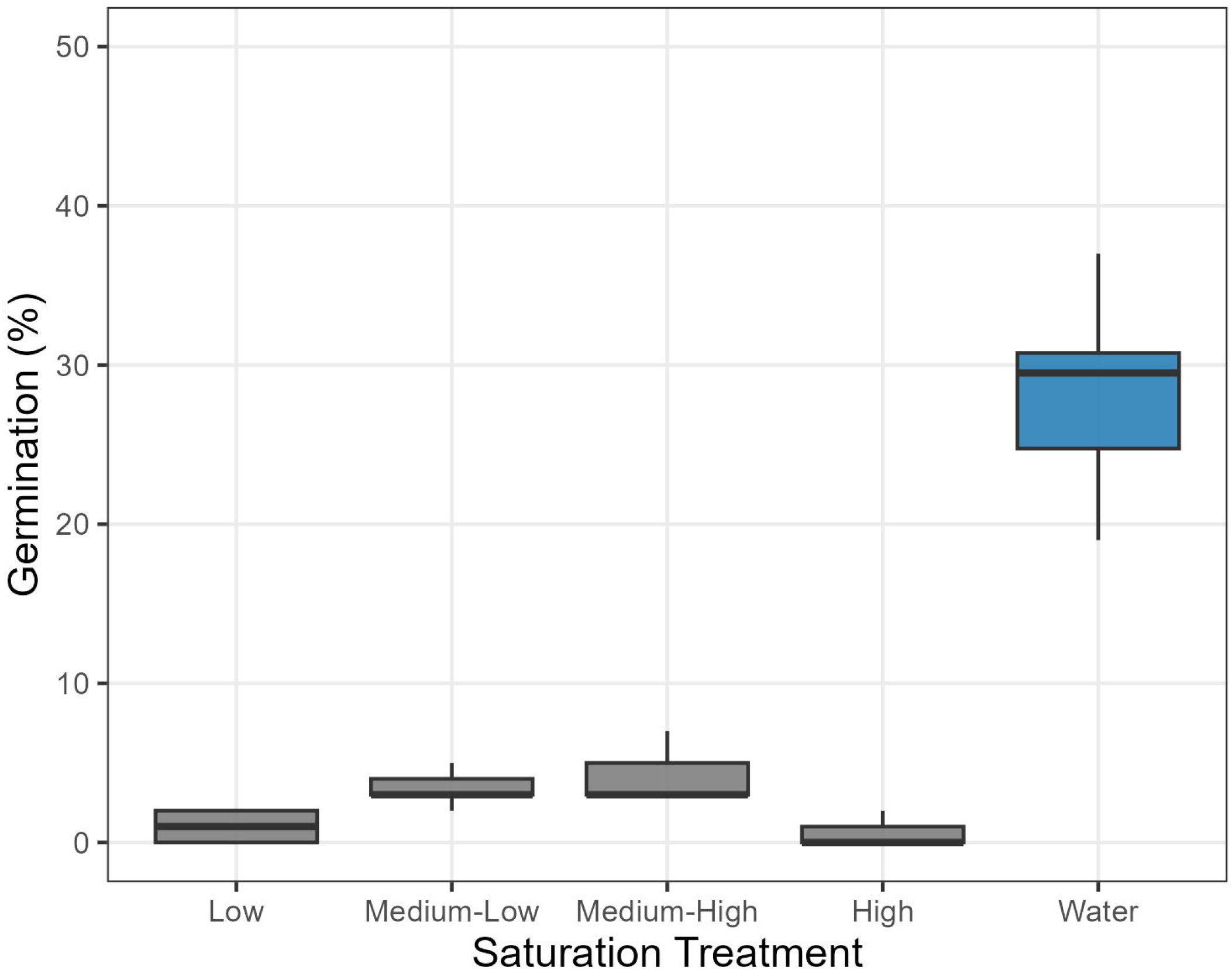
Differences in germination rates depending on the moisture conditions. Each box represents the interquartile range with the median indicated as the middle line. Outliers are not present. The low to high water treatments took place in soil, while the water treatment was submerged in pure water.

**Figure 5.**
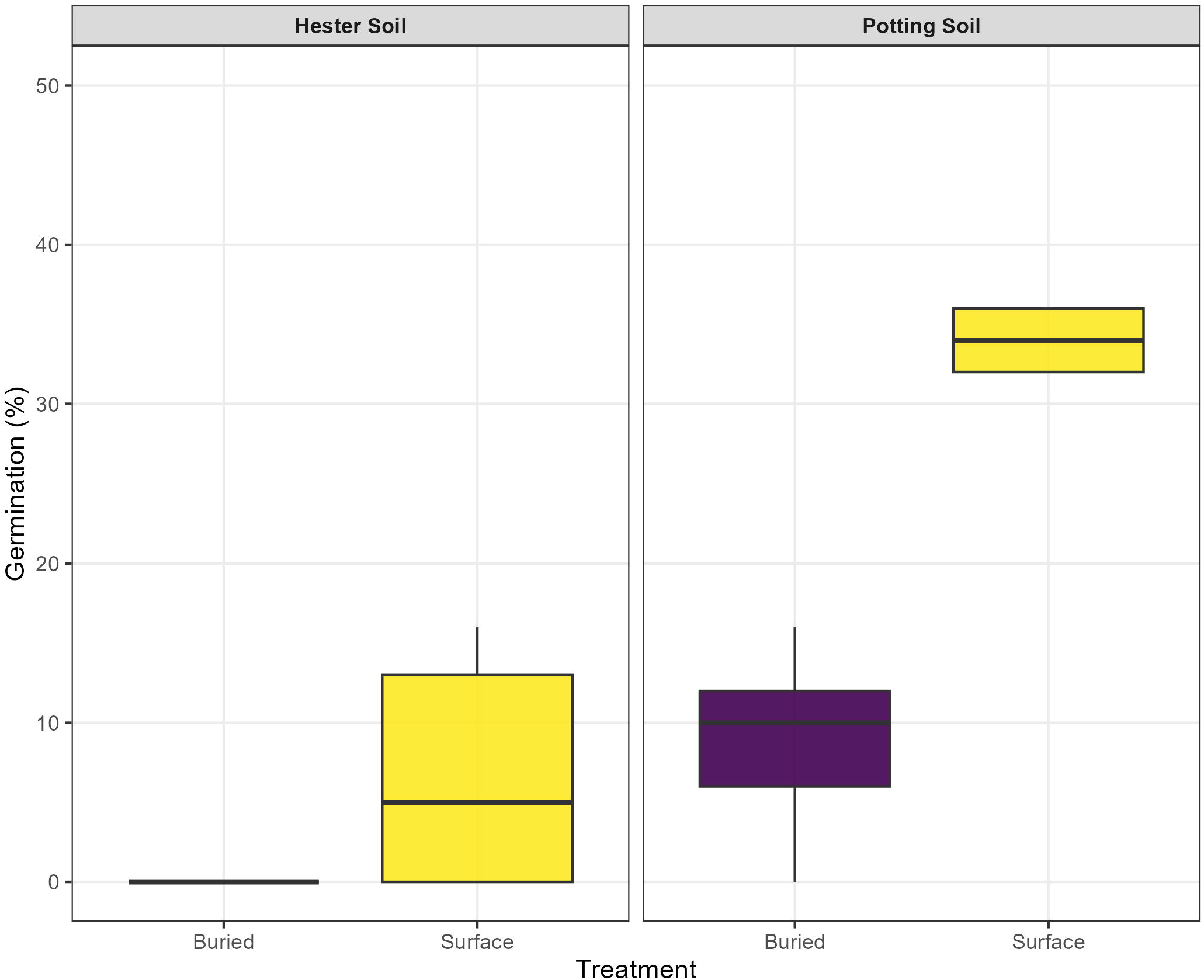
Differences in germination rates under different soil and pickleweed seed sowing conditions. Each box represents the interquartile range with the median indicated as the middle line. Outliers are present. Buried seeds were sown 0.5 cm deep and covered with soil, while surface seeds were not covered.

### Sowing Depth and Soil Type

There was a positive interaction between soil type and sowing depth (p = 0.00184, z = - 3.115). The proportion of seeds that germinated differed significantly among soil types (p < 0.001, z = 6,543) and by sowing depth (p < 0.001, z = 6.067), with greater germination observed in the seeds sown half a centimeter deep. Buried seeds in the Hester soil germinated at 0.6%, while more surface-sown seeds germinated at 6.4%; in potting soil, the percentages were 8.9 and 34%, respectively (Table 5).

## Discussion

Our investigation serves as a model for how experimental studies can help increase the success of seed-based restoration in salt marsh ecosystems. Seed additions are an underutilized revegetation method in coastal restoration due to limited knowledge of seed dynamics in intertidal environments. Previous studies from southern California revealed that salinity and moisture (Noe, 2002) and nutrient availability (Moseman, 2007) shape the successful germination of many Californian salt marsh species. Our experiments expand on this foundational knowledge, more closely investigating how salinity, moisture, seed source, and organics in the soil affect germination. We identified various critical steps that affect the success of seed-based restoration (Table 1). Below, we review three of these influential steps separately. For each step of the seed addition process, we consider both the environmental influences on establishment success as well as direct interventions to be taken by restoration practitioners to enhance vegetation success. Our study serves as a model for evidence-based restoration by approaching seed additions through a step-wise process that simplifies the methodology and links experimental findings to practical restoration recommendations.

Sourcing material with high viability increases the chance for germination and subsequent establishment success for restoration sites (Table 1A). In general, higher baseline seed viability will lead to greater germination and, therefore, greater seedling establishment. We found significant differences in viability and seed size among different sites. This finding is consistent with other seed provenance studies that found effects on viability, germination, and dormancy (Ensslin *et al*., 2018). One such study on *Spartina* seeds found that different sites produced seeds with varying characteristics, including spike length, seeds per spike, and the number of fruiting culms (Xiao, Zhang and Zhu, 2009). When investigating the effects of provenance on seeds, we found a significant effect of seed size on germination percentage (Figure 1). However, selecting seeds based on size only works if there is established data on which sources produce larger seeds. Instead, the focus should be on selecting seeds with consistently higher viability and germination rates. The highest seed viability came from within the restoration site rather than from any of our natural sites. While additional research is needed to determine the exact mechanism underlying this increase, we hypothesize that since the plants within Hester are not visually as healthy as plants in our natural marshes, it may be due to stressed plants investing more in seeds than unstressed plants.

Coastal halophytes frequently display physiological dormancy, preventing germination during unfavorable tidal or saline conditions (Ungar 2003, Baskin and Baskin 2007). Seed pretreatments are critical for breaking dormancy of many seeds and increasing germination, yet treatment effectiveness depends on both the natural history of the targeted species and the environmental conditions (Table 1C). Experimental studies can identify the pretreatments specific species may require under different stressors or soil treatments, potentially mimicking the conditions wild seeds encounter before germination. To investigate the best way to break dormancy, our stratification experiments investigated how moisture, salinity, and temperature conditions might mimic various natural events - tides or freshwater precipitation - and promote germination responses. We demonstrated that prolonged soaking significantly enhanced germination rates (Figure S6). However, responses differed depending on the soils in which the seeds were sown, indicating context-dependent germination for our seeds. Our findings are consistent with earlier studies showing high germination of marsh species following heavy rains, which would provide sustained moisture comparable to soaking (Noe and Zedler 2001, Callaway and Zedler 2004). Cold stratification replicates the temperature conditions and moisture imbibition that seeds naturally experience when they are swept away from — and then brought back to — the marsh platform by the tides. Additionally, cold stratification is a common dormancy-breaking mechanism for seeds with physiological dormancy (Baskin and Baskin, 2004). However, our stratification experiment also displayed that the optimal pretreatment may vary with the soil conditions in which the seeds are being sown.

It is critical to match seed treatments to the site-specific environmental conditions, for both dormancy-breaking pretreatments and for sowing seeds in the field (Table 1D). The conditions of restoration sites can often differ from those of natural ecosystems. For restored marshes resulting from soil additions, we might expect more soils with little to no existing organic matter. Our experiments took place in both benign soils that may be more typical of natural, highly organic marshes, as well as more stressful soils that may mimic those resulting from restoration methods. The different soil conditions led to significant differences in germination depending on the stratification pretreatment seeds experienced before being sown. Freshwater soaking (mimicking freshwater precipitation) increased germination under stressful soil conditions, while longer saltwater soaks (mimicking tidal inundation) increased germination under non-stressful conditions. The type of soil may also affect the extent to which moisture and salinity hinder or promote germination. We observed strong interactions between moisture and salinity, with intermediate moisture levels maximizing germination at lower salinity levels. Soil type further enhances these effects; seeds in potting soil exhibited faster germination and higher total emergence than those in the denser Hester mimic. This context-dependence has been documented for a variety of crops, where stratification treatments differ depending on whether the crop is being planted in an agricultural or natural setting (Catford *et al*., 2022). Such context-dependence underscores that there is no universal ‘best’ sowing condition and that success may depend on matching seeds to the local environmental conditions.

How deep seeds are sown into the soil can also determine both their ability to germinate and their vulnerability to environmental stressors. After determining how the soil conditions of the environment may adapt to different pretreatments, it is then important to consider the more technical aspects of sowing — namely, how deep to sow seeds and how much moisture they need. Our sowing depth experiment demonstrated that sowing half a centimeter deep improved emergence, particularly in looser potting soils, aligning with findings from Baldwin et al. (1996) and Zedler (2003). Many coastal species rely on light exposure to signal favorable germination windows, meaning surface sowing often yields higher germination than burial under compact or saturated soils (Martínez, Valverde and Moreno-Casasola, 1992). It is also important to consider the timing and moisture conditions of the seeds when sowing. Our moisture and submersion experiment found the highest germination when seeds remained fully submerged. This can be mimicked in natural conditions by adding seeds before or during precipitation/rainfall events. Adding seeds during rainfall events can enhance germination, but at the risk of seed loss from erosion or tidal scouring. Thus, shallow seed burial may be recommended for maintaining seeds at the locations where they were sown. Careful control of sowing depth can mimic natural dispersal while minimizing seed loss.

Active management can substantially improve seed addition outcomes by aligning restoration practices with seed biology and environmental context. Each stage of sowing seeds into coastal soils (Table 1) —sourcing, pretreatment, sowing, and establishment—requires tailored interventions informed by seed ecology. From our experimental studies, we have four recommendations for seed-based coastal restoration: (1) choose seeds from high-viability sources, (2) apply pretreatments that mimic natural dormancy cues, (3) consider the importance of context-dependent outcomes, and (4) optimize sowing depth and timing. Seed lots should be tested for baseline viability and, when possible, collected from local populations adapted to site-specific salinity and moisture regimes. Soaking or cold stratification should replicate natural conditions, which for coastal plants involves tidal inundation or seasonal flooding cycles. Sowing should coincide with seasonal windows of favorable conditions, which for these marsh plants consist of reduced salinity and high moisture availability to enhance germination. Finally, it is crucial to tailor approaches to the target restoration site and its specific stressors and soil conditions at all stages of the seed addition process. These lessons are critical to informing SBR of pickleweed and other coastal plants. More broadly, for any restoration project, management strategies that account for context-dependence will better overcome the challenges of seed dormancy and germination inhibition, and therefore enhance restoration success.

## Supporting Information

**°Figure S1. Locations of the ten sites where seeds were sourced**

**°Figure S2. Table showing differences in soil composition for the three different soil treatments**

**°Figure S3. Experimental layout of the soaking experiment**

**°Figure S4. Seed size distributions of the 10 sources. Numbers at the top represent the average size (in mm) for each site. Each box represents the interquartile range with the median indicated as the middle line. Whiskers extend to 1.5 times the interquartile range**

**°Figure S5. Seed size to germination relationship**

**°Figure S6. Days to germinate in the stratification experiment. MGT = Mean Germination Time. Control seeds were not soaked and were directly sown from storage into the soil**.

**°Figure S7. Days to germinate in the moisture/salinity inhibition experiment. MGT = Mean Germination Time. Freshwater additions were at 0ppt, while saltwater additions were 30ppt. High moisture levels were around 96% Volumetric Water Content (VWC), medium moisture levels were around 63% VWC, and low moisture levels were around 35% VWC. These reported moisture levels represent the average VWC immediately following daily watering in the greenhouse**.

**°Figure S8. Mean observed and model-estimated germination of Salicornia *pacifica* seeds under five moisture treatments**.

## Notes

<u>Financial Support:</u> Funding for travel and for field monitoring by Reserve staff was provided by various grants supporting Elkhorn Slough’s Hester Marsh restoration, including California Ocean Protection Council and NOAA Climate Program Office’s National Integrated Drought Information System (ROR https://ror.org/00mmmy130) under award NA22OAR4310221-T1-01.

### Competing Interest Statement

The authors have declared no competing interest.

