## Supplementary figures and images for "Factors Affecting Germination of a Dominant Salt Marsh Species are Context-Dependent: Implications for Coastal Seed-Based Restoration"

### Supplemental Figure 1

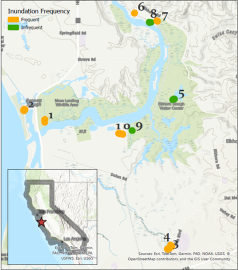

### Supplemental Figure 2

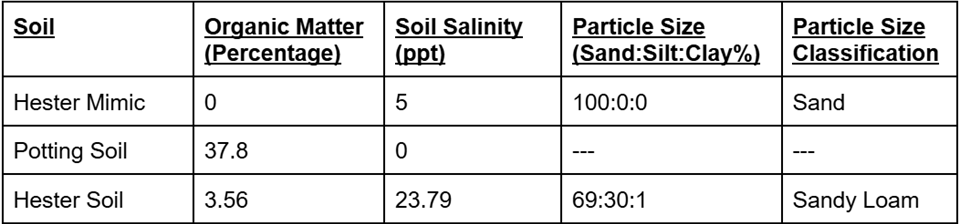

### Supplemental Figure 3

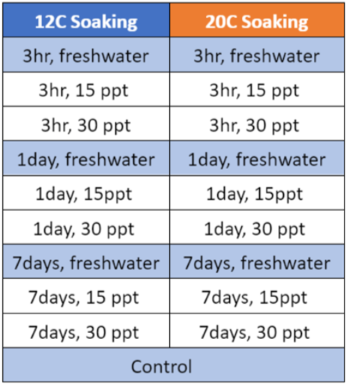

### Supplemental Figure 4

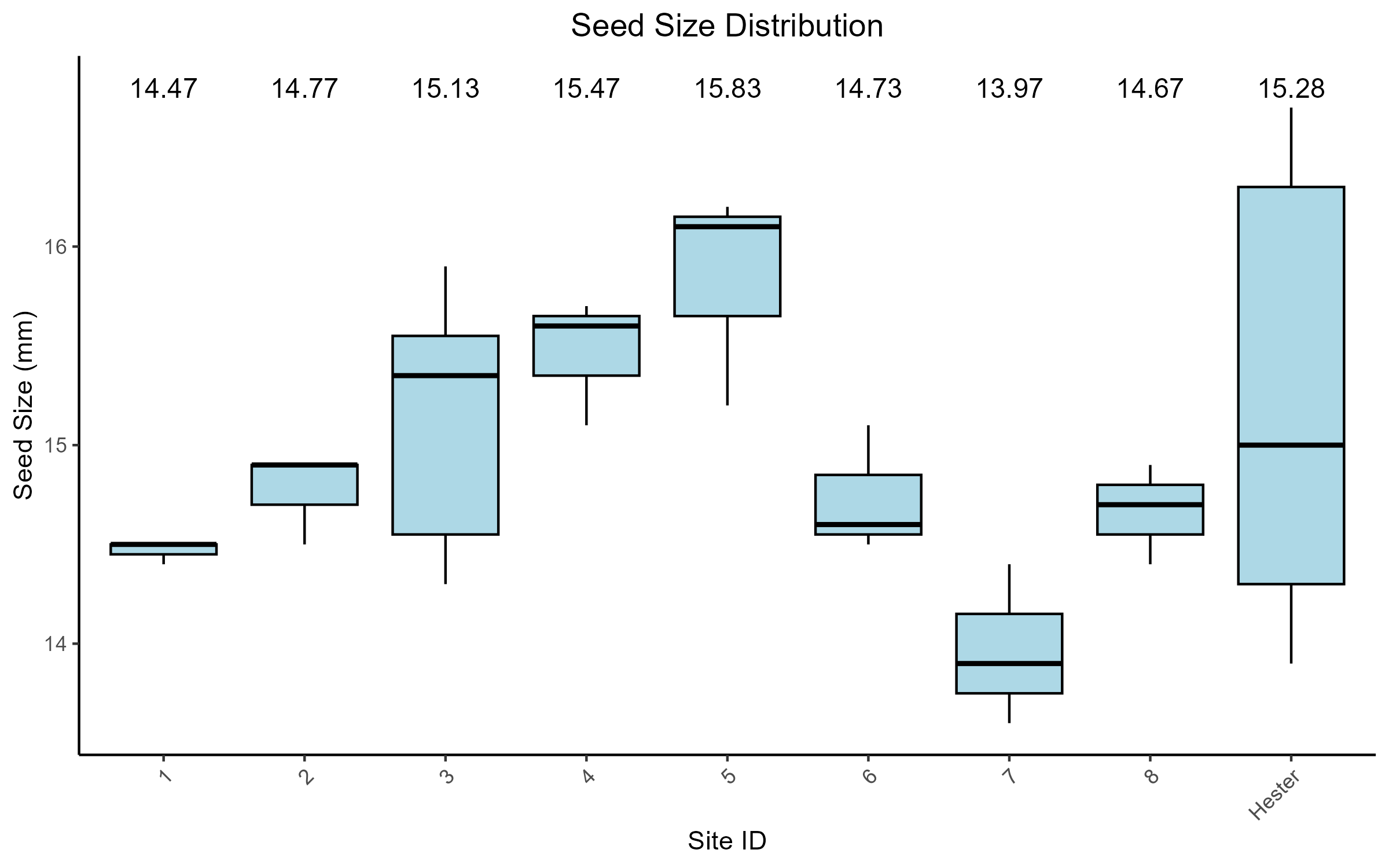

### Supplemental Figure 5

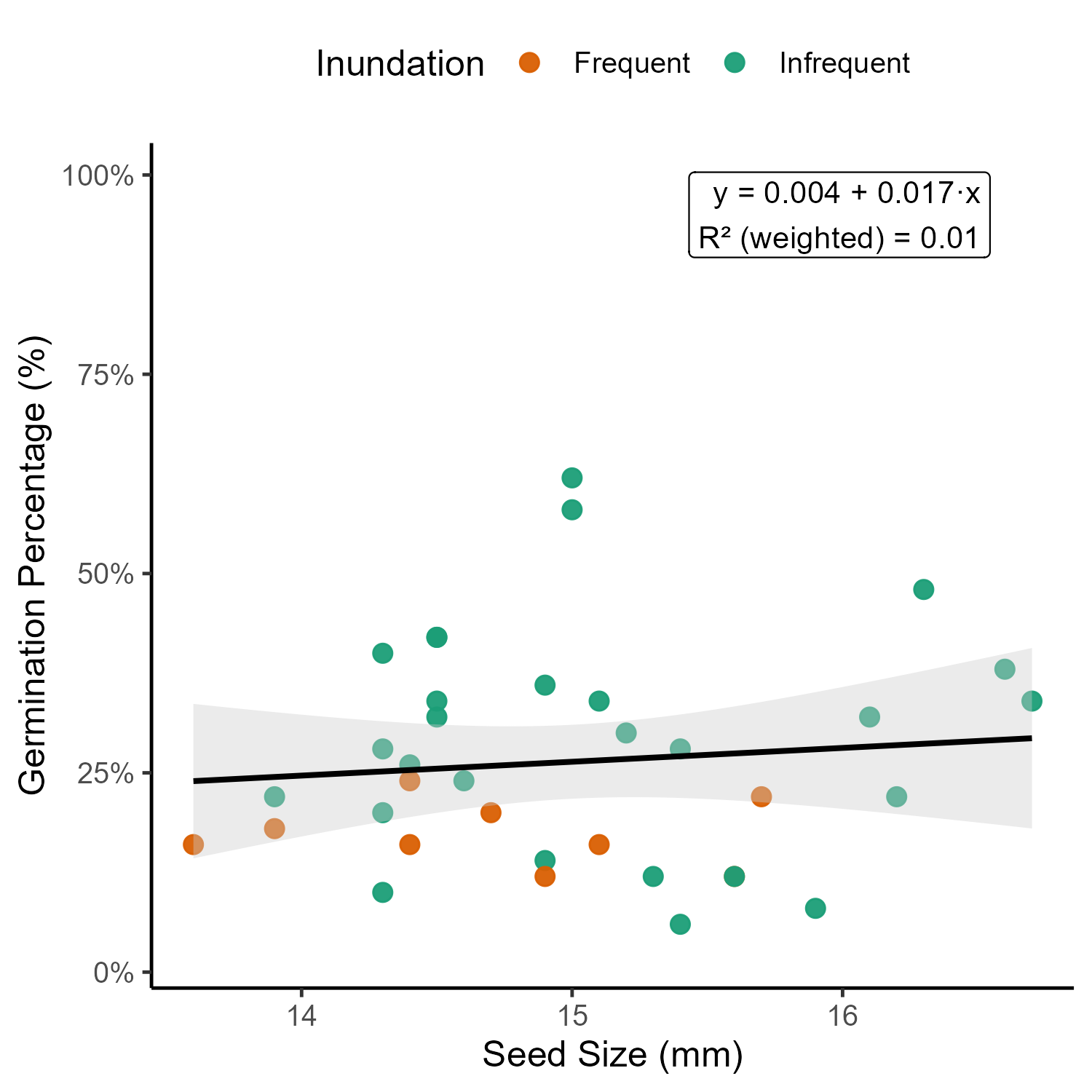

### Supplemental Figure 6

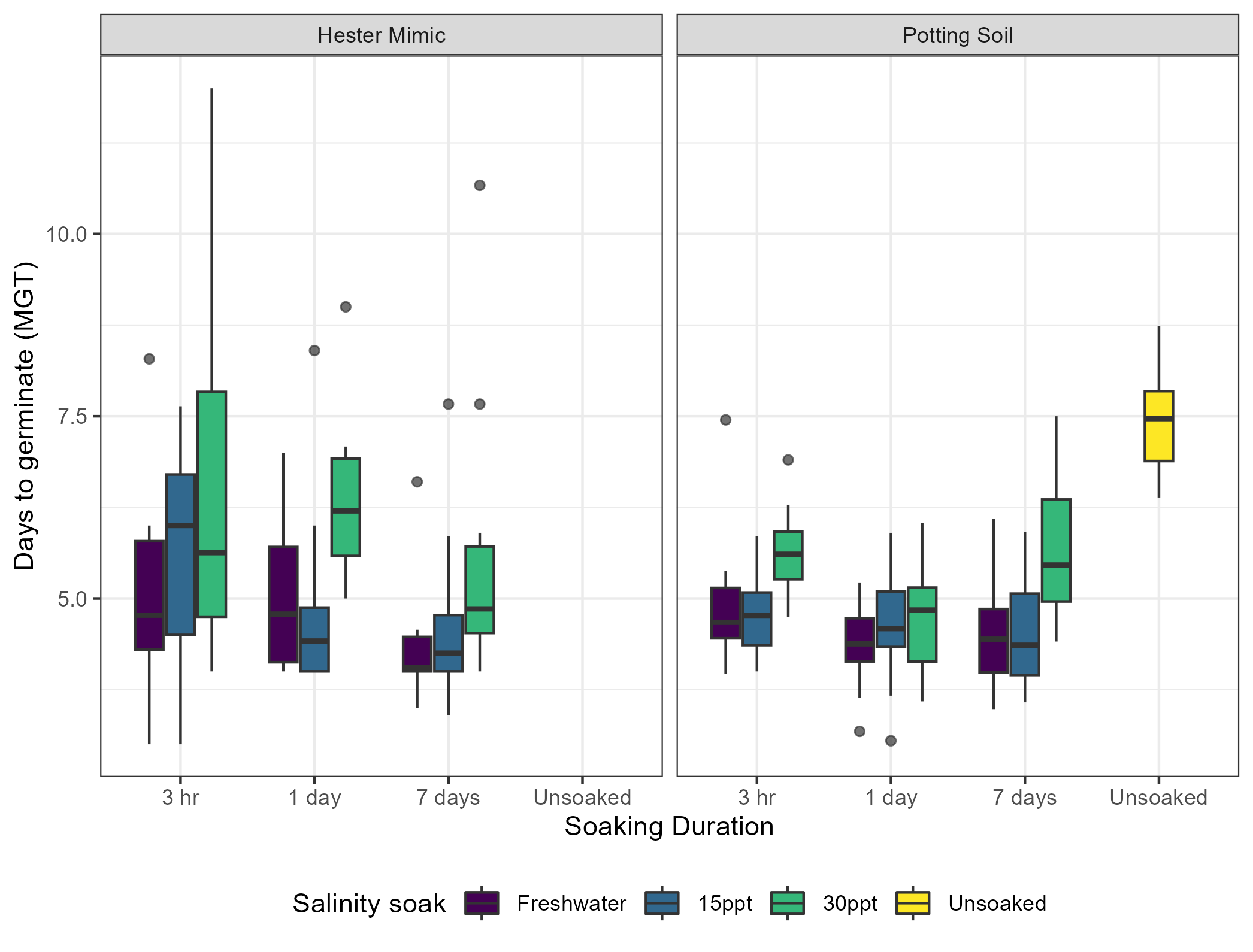

### Supplemental Figure 7

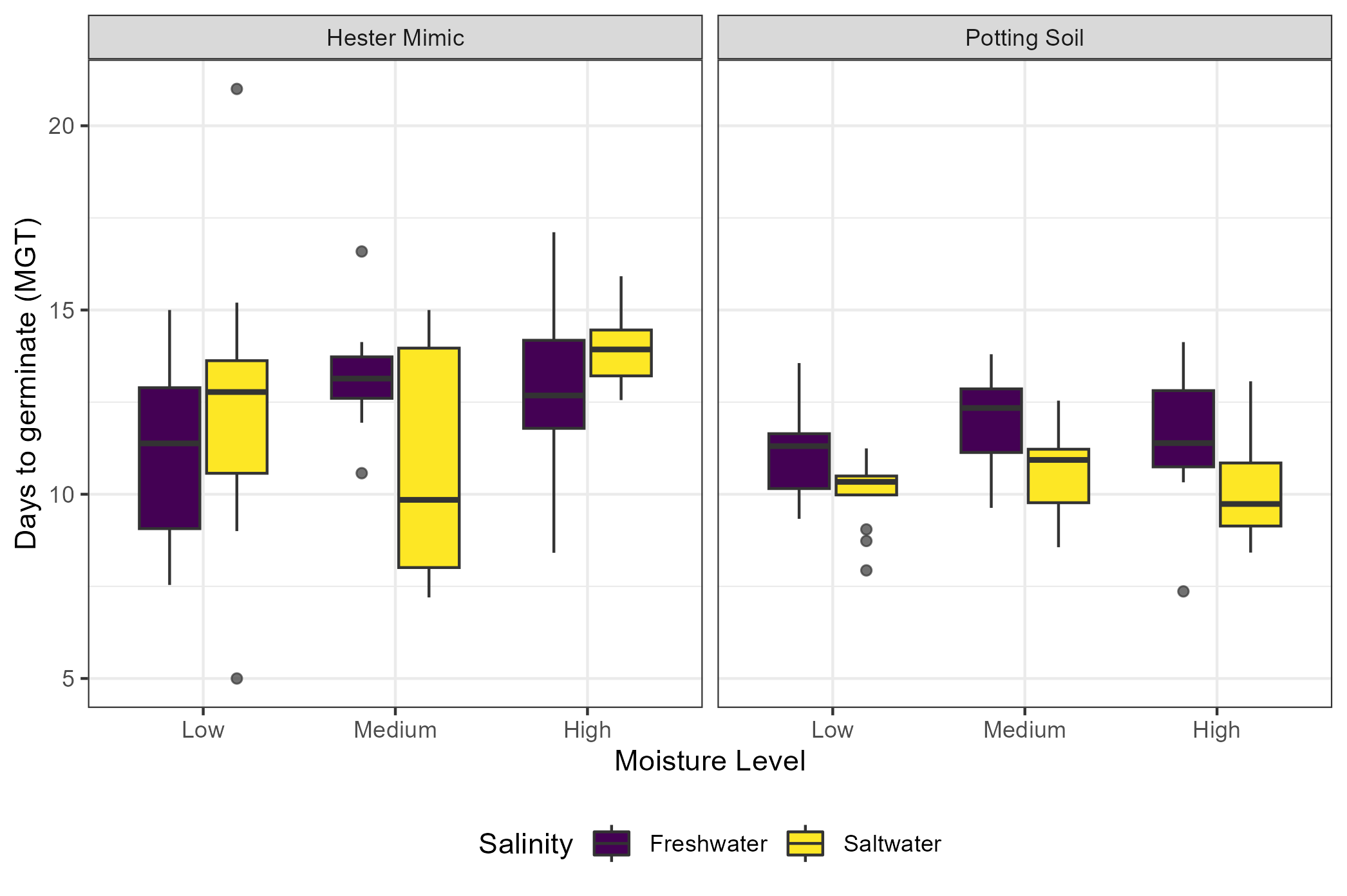

### Supplemental Figure 8

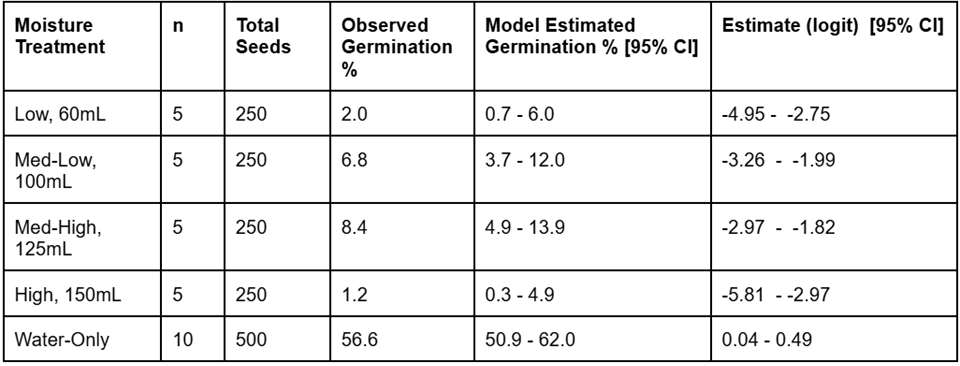
